# Plasticity, repeatability, and phenotypic correlations of aerobic metabolic traits in a small estuarine fish

**DOI:** 10.1101/2020.05.01.072587

**Authors:** Jessica E. Reemeyer, Bernard B. Rees

**Author notes:** McGill University, Department of Biology, 1205 Avenue Dr Penfield, Montreal, Quebec, Canada, H3A 1B1 (Jessica E. Reemeyer). Corresponding Author: Jessica E. Reemeyer, McGill University, Department of Biology, 1205 Avenue Dr Penfield, Montreal, Quebec, Canada, H3A 1B1.

## Abstract

Standard metabolic rate (SMR), maximum metabolic rate (MMR), absolute aerobic scope (AAS), and critical oxygen tension (P_crit_) were determined for the Gulf killifish, *Fundulus grandis*, an ecologically dominant estuarine fish, acclimated to lowered salinity, elevated temperature, and lowered oxygen concentration. Acclimation to low salinity resulted in a small, but significant, elevation of P_crit_; acclimation to elevated temperature increased SMR, MMR, AAS, and P_crit_; acclimation to low oxygen led to a small increase in SMR, but substantial decreases in MMR, AAS, and P_crit_. Variation in these metabolic traits among individuals was consistent and repeatable when measured during multiple control exposures over seven months. Trait repeatability was unaffected by acclimation condition suggesting that repeatability of these traits is not context dependent. There were significant phenotypic correlations between specific metabolic traits: SMR was positively correlated with MMR and P_crit_; MMR was positively correlated with AAS; and AAS was negatively correlated with P_crit_. In general, within-individual variation contributed more than among-individual variation to these phenotypic correlations. The effects of acclimation on these traits demonstrate that aerobic metabolism is plastic and influenced by the conditions experienced by these fish in the dynamic habitats in which they occur; however, the repeatability of these traits and the correlations among them suggest that these traits change in ways that maintains the rank order of performance among individuals across a range of environmental variation.

**SUMMARY STATEMENT:** Aerobic metabolism of an ecologically dominant estuarine fish is influenced by acclimation to environmental changes without altering trait repeatability. Furthermore, specific metabolic traits are phenotypically correlated.

## INTRODUCTION

Elucidating the causes and consequences of variation in energy metabolism is a central goal of animal physiology and ecology (Schmidt-Nielsen, 1997; Brown et al., 2004). In particular, there is considerable interest in the effects of changes in the abiotic environment on the intensity of energy metabolism, an interest that has been heightened by the rate and extent of contemporary alterations in climate and other environmental variables brought about by human activities. Estuaries are naturally dynamic habitats due to variable riverine input, diurnal and tidal cycles, wind patterns, storms, and season. As a result, several abiotic factors, including salinity, temperature, and dissolved oxygen (DO), vary broadly on time scales ranging from hours to months. Anthropogenic changes in land use, hydrology, and climate can alter the mean value of these abiotic variables (e.g., higher average temperature) or increase the frequency and duration of extreme values (e.g., aquatic hypoxia). Understanding the response of energy metabolism in estuarine organisms to natural environmental variation will provide insight into their resiliency and potential responses to future habitat alteration.

The metabolic rate of an animal is an integrative measure of energy flow and includes costs of ion transport, biosynthesis, locomotion, and myriad other processes. For organisms that utilize oxidative phosphorylation for ATP synthesis, metabolic rate can be estimated as the rate of oxygen consumption (M_O2_). In ectothermic animals, including fishes, the M_O2_ in a post-absorptive individual at rest is its standard metabolic rate (SMR), while maximum metabolic rate (MMR) is the highest M_O2_, typically achieved during or immediately after intense exercise (Brett and Groves, 1979; Norin and Clark, 2016). The difference between MMR and SMR is absolute aerobic scope (AAS), which represents the animal’s capacity to support energetically expensive processes aerobically. At low levels of ambient oxygen (hypoxia) aerobic metabolism (SMR, MMR, and AAS) becomes limited (Fry, 1947). The level of oxygen below which SMR can no longer be sustained is the critical oxygen tension, or P_crit_ (Rogers et al., 2016; Claireaux and Chabot, 2016; Reemeyer and Rees, 2019). Below P_crit_, metabolism is increasingly supported by anaerobic processes (e.g., glycolysis), which are not sustainable over the long term, or it is reduced through metabolic suppression (Richards, 2009). Animals that can maintain SMR down to lower levels of oxygen have lower P_crit_; therefore, P_crit_ has been proposed as an index of hypoxia tolerance (Rogers et al., 2016).

In fishes, SMR, MMR, AAS, and P_crit_, are influenced by changes in salinity, temperature, and oxygen. Bony fish regulate the ion composition of plasma at about 1/3 that of sea water. At salinities lower and higher than this, passive fluxes of ions out of or into the fish are balanced by energy-dependent ion transport, which is predicted to increase SMR (Bœuf and Payan, 2001), although this cost may be small (Ern et al., 2014). In addition, the “osmo-regulatory compromise” posits that because passive ion fluxes occur across the same tissue responsible for respiratory gas exchange (gill), changes that serve to limit one may also limit the other (Sardella and Brauner, 2007). Thus, a reduction of gill surface area, which could reduce passive ion flux, could potentially limit MMR or increase P_crit_. As temperature increases, SMR and MMR generally increase; however, SMR increases exponentially, while MMR reaches a plateau or decreases at high temperatures (Pörtner, 2010; Pörtner and Farrell, 2008). The effect of temperature on AAS, therefore, is for it to increase at moderate temperatures, but then decline. At high temperatures, AAS may fall to zero if SMR equals MMR (Pörtner, 2010; Pörtner and Farrell, 2008). P_crit_ is also affected by temperature, increasing at higher temperatures (Rogers et al., 2016; McBryan et al 2016). Exposure to low oxygen limits aerobic metabolism, especially MMR and AAS, as the oxygen available to support activities beyond SMR decreases. Acclimation of fishes to hypoxia, however, may improve oxygen extraction, which has been documented as a decrease in P_crit_ (Borowiec et al., 2015).

There is growing appreciation that traits related to aerobic metabolism are repeatable when measured in the same individuals over time. As pointed out by Bennett (1987) and more recently by others (Roche et al., 2016; Killen et al., 2016a), repeatable individual variation in physiology arises from differences in genetics, development, and environment. When the variation is due to genetic factors and is heritable, it represents the raw material upon which natural selection can act. SMR, MMR, and AAS have been shown to be significantly repeatable across multiple measures among individuals of several species (Killen et al., 2016a; Nespolo and Franco, 2007; Norin and Malte, 2011; Norin et al., 2016; Virani and Rees, 2000). Although there are fewer reports of the repeatability of P_crit_, it, too, appears to be a repeatable trait (Pan et al., 2018; Reemeyer and Rees, 2019). Currently, however, there is a lack of studies on whether the repeatability of aerobic metabolism is altered by acclimation to changes in abiotic variables. Norin et al. (2016) showed that acute exposure of Barramundi (*Lates calcarifer*) to low salinity, elevated temperature, and low oxygen differentially affected individuals with low or high metabolic rate. The result was that the rankings of individuals’ SMR, MMR, and AAS were different during acute exposure than during control conditions. Because that study examined only short-term acute exposures, it remains to be seen how acclimation to these conditions affects repeatability of these traits.

An experimental framework to assess repeatability of traits can also be valuable in illuminating the relationships between traits. Traditionally, studies of the relationship between traits measure each trait once each in a group of organisms. A repeated-measured design, where each trait is measured multiple times in each individual, enables the determination of covariance in the traits both among- and within-individuals (Dingemanse and Dotcherman, 2013; Careau and Wilson, 2017). The phenotypic correlations (r_p_) can then be partitioned into among-individual correlations (r_ind_) and within-individual correlations (r_e_); with r_ind_ representing linkages in traits due to a combination of genetic and fixed environmental factors, and r_e_ representing linkages due to a combination of shared plasticity and correlated measurement error (Brommer, 2013; Careau et al., 2014). High estimates of r_ind_ indicate that across individuals, those with high values for one trait also have high values for another trait (and *vice versa*); whereas, when r_e_ is high, the interpretation is that for a given individual measured at a specific point in time, the values for the two traits are high, but when measured at another point in time, both traits could be low.

In the present study, SMR, MMR, AAS, and P_crit_ were measured in the Gulf killifish, *Fundulus grandis*, after acclimation to lowered salinity, elevated temperature, and lowered DO. *Fundulus grandis* is a small-bodied, abundant species found throughout estuaries along the Gulf of Mexico (Nordlie, 2006). Because they inhabit dynamic environments and tolerate large fluctuations in abiotic variables, *F. grandis* and its sister species *F. heteroclitus* are excellent model species for environmental biology research (Burnett et al., 2007). Fish used in this study were collected at the Grand Bay National Estuarine Research Reserve (GBNERR), part of a system of protected areas in coastal U.S.A. Specifically, *F. grandis* were collected at two sites within the GBNERR, which long-term water quality records indicated were similar in salinity and temperature, but differed in annual DO profiles. This allowed evaluation of local differences in aerobic metabolism, as well as potential differences in the effects of acclimation to low oxygen. The goals of this study were to address the following questions: (1) How are SMR, MMR, AAS, and P_crit_ affected by acclimation to low salinity, high temperature, and low oxygen? (2) What are the long-term repeatability estimates of SMR, MMR, AAS, and P_crit_ in *F. grandis*? (3) Does acclimation to low salinity, elevated temperature, and low oxygen affect the repeatability of SMR, MMR, AAS, and P_crit_? (4) Do individuals from different collection sites respond differently to low salinity, elevated temperature, and low oxygen? (5) Are there phenotypic correlations between these traits, and if so, do these arise from among-individual correlations, within-individual correlations, or both.

## MATERIALS AND METHODS

### Field sites and fish collection

*Fundulus grandis* were collected at the GBNERR in August 2018. The GBNERR is a protected wetland within the Mississippi Coastal Streams Basin (Fig. S1) and includes several sites with long-term water quality data. Among these are Bayou Heron (BH, 30.4178° N, 88.4054° W) and Bayou Cumbest (BC, 30.3836° N, 88.4364° W), which have comparable salinity and temperature, but differ in DO profiles (Fig. 1). Bayou Heron experiences summer hypoxia due to enhanced nutrient input from natural sources, while Bayou Cumbest remains normoxic throughout the year. Approximately equal numbers of fish (50 each) were collected at the two sites using baited minnow traps placed along the marsh edge within 1 km of permanently moored water quality sensors, or sondes (Fig. S1).

**Figure 1:**
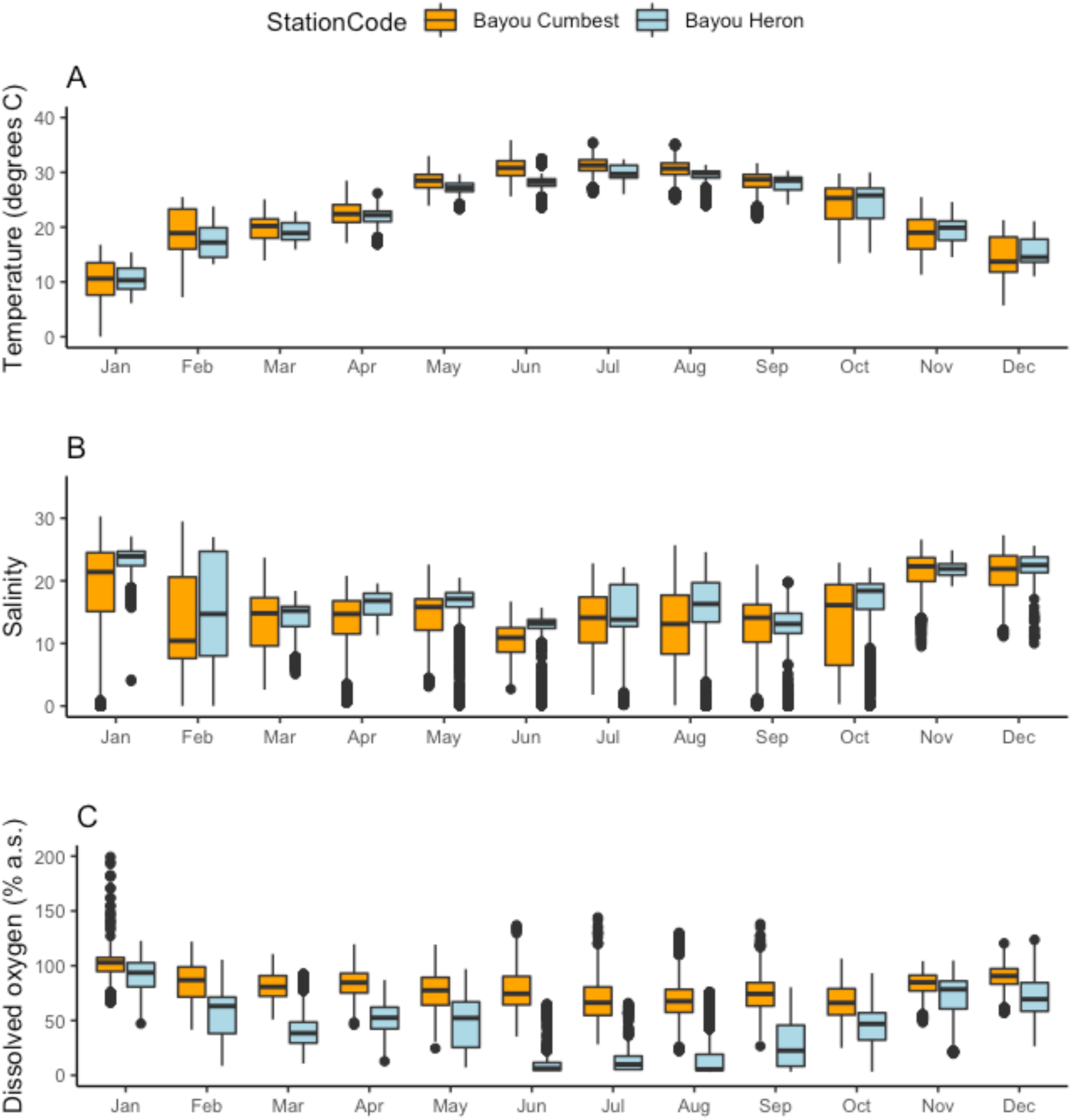
Annual variation in water temperature (A), salinity (B), and DO (C) at the Bayou Cumbest (orange) and Bayou Heron (blue) monitoring sondes. Data were collected every 15 min and compiled for 1 August 2017 to 31 September 2018. Box and whisker graphs show medians (center line), upper and lower quartiles (box), and total data range (whiskers) after removing outliers (black dots).

To account for fine scale variation in water quality, temperature, salinity, and DO were measured with a hand-held meter (YSI Pro2030, www.ysi.com) at the exact times and locations of trap deployment and retrieval (Table S1). These measurements showed that temperature was essentially identical between collection sites, as well as between sondes and hand-held measurements, although salinity was lower at the trap locations which were more affected by freshwater runoff from recent rains. In support of long-term records, the average and range of DO were lower at the BH sites than the BC sites, but this difference was not as pronounced as the difference recorded over the same period by the sondes. This discrepancy was attributed to the fact that sondes were located 1-2 m from the surface, and thus more influenced by vertical stratification of the water column than the trap sites, which were all in the top 0.5 m of the water column.

### Fish husbandry

Fish were held at the GBNERR for up to 2 days and then transported to the University of New Orleans in aerated field-collected water. All individuals were treated prophylactically for external parasites using API General Cure (www.apifishcare.com) according to the manufacturer’s instructions within 1 week of collection. Fish were maintained for at least 6 weeks in 38 l aquaria containing aerated, filtered, dechlorinated tap water adjusted to salinity ≈ 10 using Instant Ocean Synthetic Sea Salt (www.instantocean.com). The photoperiod was 12:12 (light:dark) and temperature was approximately 25°C. During the initial 6-week period, fish were fed twice daily to satiation with Tetramarine Large Saltwater Flakes (www.tetra-fish.com). Thereafter, fish were fed 1-1.5% of the total fish mass once per day, except for the days prior to and including respirometry (see below). After 6 weeks, fish were tagged with passive integrated transponder (PIT) tags according to Reemeyer et al. (2019). Fish were maintained for a minimum of 1 week after tagging before experiments. All fish maintenance and experimental procedures were approved by the University of New Orleans Institutional Animal Care and Use Committee (protocol no. 18-006).

### Acclimation regimes and morphological measurements

One week prior to the experiment, two groups of 30 fish were moved into one of two 100 l tanks and allowed to adjust to the new tanks. Fish were selected to achieve roughly equal numbers of males and females from both BH and BC, and they ranged in size from 2 to 6 g. The experiment consisted of serially acclimating fish over a period of approximately 7 months across a range of salinity, temperature, and DO similar to those determined at the time of collection. Each exposure interval lasted 4 weeks and followed the order (1) control conditions (T = 25°C, salinity = 10, DO > 85% a.s., 6.6 mg l^-1^, 17.5 kPa), (2) low salinity (salinity = 1), (3) control conditions, (4) high temperature (T = 32°C), (5) control conditions, (6) low oxygen (DO = 30% a.s., 2.3 mg l^-1^, 6.2 kPa), and (7) control conditions. For each interval, water was adjusted to the desired conditions on day 1 as described below. Fish were then held under these conditions for 14 d, respirometry was conducted over the next 12 d, and fish recovered 1 d prior to the next change in conditions. Over the 12-d respirometry period, batches of 4 fish were randomly selected for measurement (see below). Consequently, the acclimation period prior to respirometry ranged from 15 to 26 d. Importantly, the results reported below include the effects of acclimation as well as measurement under the same conditions. Water quality in the two acclimation tanks did not differ, and the mean and range of temperature, salinity, and DO during each interval are shown in Table S2.

For the low salinity acclimation, salinity was lowered from 10 to 1 over the course of 9 h by gradual replacement of tank water with dechlorinated tap water. At the end of the low salinity treatment, salinity was raised over the course of 9 h by adding artificial sea salt to achieve a final salinity of 10. For the high temperature acclimation, temperature was increased from 25°C to 32°C over the course of 7 h at a rate of 1°C per h using a digital heater controller connected to a titanium aquarium heater (www.Finnex.com). Temperature was maintained using the same controller and heater over the course of the experiment. At the end of the high temperature exposure, water temperature was lowered by turning down the heater and replacing part of the tank water with water at 25°C. For the low oxygen acclimation, DO was lowered at approximately 10% a.s. per h over 7 h by gassing tank water with nitrogen. DO was continuously monitored by a galvanic oxygen sensor (www.atlas-scientific.com) connected to a raspberry pi computer (www.raspberrypi.org). The computer was programmed to take input from the oxygen sensor once per min and control the introduction of nitrogen from a gas cylinder via a solenoid valve to achieve the desired DO level. At the end of hypoxic acclimation, nitrogen introduction was halted, and water was aerated with aquarium air pumps to achieve an increase of approximately 10% a.s. per h until DO exceeded 85% a.s.

At the conclusion of each interval, and before beginning the next exposure, all fish were lightly anaesthetized in dechlorinated, salinity-adjusted water with 0.1 g l^-1^ MS-222, gently blotted, measured for mass (M) and standard length (SL). Fulton’s (1904) condition factor (K) was calculated as M SL^-3^. Daily specific growth rate (SGR) was calculated as 100 (e^G^ – 1) where G = (ln(M_2_) – ln(M_1_))(t_2_ – t_1_)^-1^, where M_1_ and M_2_ are masses determined at two times, t_1_ and t_2_ (Stierhoff et al., 2003). Over the course of the 7-month experiment, fish increased in M and SL (Table S3). In general, SGR was low (0.03 – 0.07% body mass d^-1^) but increased during acclimation to 32°C (0.47 ± 0.20% d^-1^) and the subsequent control interval (0.23 ± 0.18% d^-1^). Condition factor (K) remained relatively constant, and fish appeared to be in good health throughout the experiment.

### Respirometry

Intermittent-flow respirometry was used to measure oxygen consumption rates (M_O2_) as described by Svendsen et al. (2016) and Reemeyer et al. (2019). The respirometry system consisted of four respirometers, each having a cylindrical glass chamber of either 118 ml or 245 ml, chosen to maintain a ratio of chamber volume to fish mass between 20 and 50 (Svendsen et al., 2016). Each chamber was fitted with two sets of tubing. One set of tubing formed a loop with a water pump that continuously circulated water from the chamber past an optical oxygen sensor and back to the chamber. The oxygen sensor was connected to a Witrox-4 oxygen meter and the oxygen saturation was measured once per second using AutoResp software (Loligo Systems; www.loligosystems.com). The second set of tubing was connected to a second pump that intermittently flushed the chamber with water from a surrounding reservoir (ca. 10 l). Water was shared among the four reservoirs (one for each respirometer) and was the same salinity, temperature, and oxygen level as the acclimation condition. This water was continuously circulated through a UV-sterilizer and heat exchanger, which, along with small aquarium heaters in each reservoir, maintained water temperature within 0.1°C of the target temperature (either 25 or 32°C). The flushing water pumps and heaters were connected to a DAQ-M relay system (Loligo Systems; www.loligosystems.com) and controlled by AutoResp software. During measurements at low oxygen, the DO in reservoir water was controlled with an apparatus identical to the one controlling DO during low oxygen acclimation.

Fish were fasted 24 h prior to respirometry. For each trial, MMR, SMR, and P_crit_ were determined sequentially over approximately 20 h. Between 15:00-16:00 fish were weighed (to the nearest 0.01 g) and placed into a circular arena (diameter = 55 cm) filled with approximately 8 l of water and chased by hand for 3 min to induce exhaustion (see pilot studies below). Immediately following the chase protocol fish were placed into the respirometer, and a cycle of 60 s flush, 30 s wait, and 120 s M_O2_ measurement was started. After 1 h, the cycle was adjusted to 300 s flush, 60 s wait, and 240 s M_O2_ measurement, which was continued for approximately 17 h. Throughout the combined ∼18 h period, *P*_O2_ was maintained at > 80% a.s., except for measurements during acclimation to low oxygen, when it was ∼30% a.s.. At 10:00 the following morning, the flush pumps were turned off, thereby creating a closed system (respirometry chamber, tubing, recirculating pump, and oxygen sensor), after which the *P*_O2_ declined due to M_O2_ by the fish. During this closed period, M_O2_ was measured over consecutive 60 s intervals until there were at least five M_O2_ measurements below that individual’s SMR. The closed period generally lasted about 60 min, after which the flush pumps were turned on to reoxygenate the chambers. All fish were given at least 10 min to recover, after which they were returned to their holding tank.

Background microbial respiration in each chamber of the respirometry system was measured before and after each trial using the following settings: 300 s flush, 60 s wait, and 1200 s M_O2_ measurement. Two M_O2_ measurements immediately before each trial were averaged, and two M_O2_ measurements immediately after each trial were averaged. Then, a time-corrected value for background respiration was subtracted from fish M_O2_, assuming a linear increase in microbial respiration over the duration of the measurement period (Reemeyer et al., 2019; Rosewarne et al., 2016). When microbial respiration exceeded 0.1 μmol min^−1^, or about 25% of the mean SMR value, the entire respirometry system was drained and sanitized with dilute bleach. This corresponded to two trials under all acclimation conditions except for high temperature where microbial respiration increased more quickly and the respirometry system was sanitized after every trial. The oxygen sensors were calibrated every 2 weeks using vigorously aerated water (100% a.s.) and water deoxygenated by the addition of sodium sulfite (0% a.s.) at the salinity and temperature of the given experimental interval.

Pilot studies validated the method to determine MMR. No significant difference in MMR was found between chasing fish for a 3 min period (above) compared to a chase of 5 min or until the fish stopped responding to a tail pinch (Brennan et al., 2016; Healy and Schulte, 2012). In addition, MMR was not higher if fish were held in air for 60 s after the chase protocol, a treatment shown to yield higher MMR in other species (Norin and Clark, 2016; Roche et al., 2013). Furthermore, MMR was taken as the single highest M_O2_ estimate during the entire respirometry trial. Typically, this is assumed to occur within the first few minutes of the chase protocol (Clark et al., 2013). This was true for many trials in the present study; however, in numerous trials, the highest M_O2_ occurred several hours after the chase protocol, frequently coinciding with the room lights turning off (20:00) or on (08:00). In the present study (including over 300 respirometry trials), the median time to MMR was 4 h after chasing, and the M_O2_ measured immediately after chasing would have seriously underestimated MMR.

SMR was calculated as the 20% quantile of 60 M_O2_ measurements made during the dark phase of the photoperiod (between 20:00-06:00). This method has been advocated by others (Chabot et al., 2016) and produces reliable estimates of SMR for *F. grandis* (Reemeyer and Rees, 2019). Absolute aerobic scope (AAS) was calculated as the difference between MMR and SMR. P_crit_ was determined during the period of closed respirometry (10:00 – 11:00). Linear regression was fit to values of M_O2_ after it dropped below and remained below SMR. Based upon this relationship, P_crit_ was determined as the oxygen level at which M_O2_ equalled that fish’s SMR, determined in the immediately preceding overnight intermittent-flow respirometry trial (Claireaux and Chabot, 2016; Reemeyer and Rees, 2019).

### Statistical analyses

To address the goals of this study, it was important to measure aerobic metabolism during the maximum number experimental intervals. Out of a total of 60 fish used here, 36 were measured at all experimental intervals and an additional 7 fish were measured in 6 of the 7 intervals. Hence, data from these 43 fish were included in these analyses. All statistical calculations were performed in R v3.3.3 (R Core Team, 2017).

Univariate linear mixed models (LMMs) were fit using the lme4 package in R (Bates et al., 2015). Response variables (SMR, MMR, AAS, and P_crit_) were log_10_-transformed and then z-transformed to a mean of 0 and standard deviation of 1. Models included salinity, temperature, DO, interval number, sex, collection site, and log_10_ mass as fixed factors, and individual ID as a random (intercept) factor. Acclimation treatments (salinity, temperature, and DO) were included as categorical variables, whereas interval number was included as a continuous variable to account for any time-dependent change in response variables over the duration of the experiment (Biro and Stamps, 2015). Initially, all factors and two-way interactions were included in the models and then removed in a stepwise fashion when doing so improved model fit [judged by a decrease in the Akaike information criterion (AIC) greater than 2]. Based on this criterion, all interaction terms were removed, and the minimum adequate model and AIC are presented for each response variable.

Pearson’s correlation coefficient, r, was used to compare mass-corrected residuals of SMR, MMR, AAS, and P_crit_ between all possible pairs of intervals. Adjusted repeatabilties (R_adj_) were determined to estimate the repeatability of metabolic traits over the entire experiment (Stoffel et al., 2017). This approach uses an LMM framework that includes the effects of fixed factors, uses parametric bootstrapping to determine confidence intervals, and calculates statistical significance by likelihood ratio tests. Bootstrapping of 10,000 simulations was used in this study.

Phenotypic correlations (r_p_) were calculated and partitioned into among-individual (r_ind_) and within-individual (r_e_) correlations as outlined in Roche et al. (2016) and Houslay and Wilson (2017). Briefly, log_10_, z-transformed response variables were fit with bivariate mixed models using the MCMCglmm package in R (Hadfield 2010) with mass, salinity, temperature, DO, and interval as fixed factors, and individual as a random factor. The settings for the model fitting were: nitt = 390,000, burnin = 9000, and thin = 100. The covariance coefficients were then extracted from the models and used to calculate r_p_, r_ind_, and r_e_ using equations adapted from Dingemanse et al. (2012) as outlined in Careau and Wilson (2017). The highest posterior distribution (HPD) interval was calculated for each estimate as a measure of credibility, analogous to the 95% confidence interval used in frequentist statistics.

Data from this study will be made freely available on figshare.com upon acceptance of this manuscript.

## RESULTS

### Mass effects on aerobic metabolism

SMR, MMR, and AAS were positively related to body mass at all control and acclimation intervals (Table 1; Figs. S2-S5). When expressed as the relationship, M_O2_ = aM^b^, values for the scaling coefficient, b, ranged from 0.73 to 1.03 for SMR, from 0.98 to 1.34 for MMR, and from 0.98 to 1.49 for AAS. On the other hand, P_crit_ was negatively related to mass (Table 1; Fig. S5), with scaling coefficients from -0.18 to -0.41.

**Table 1:**
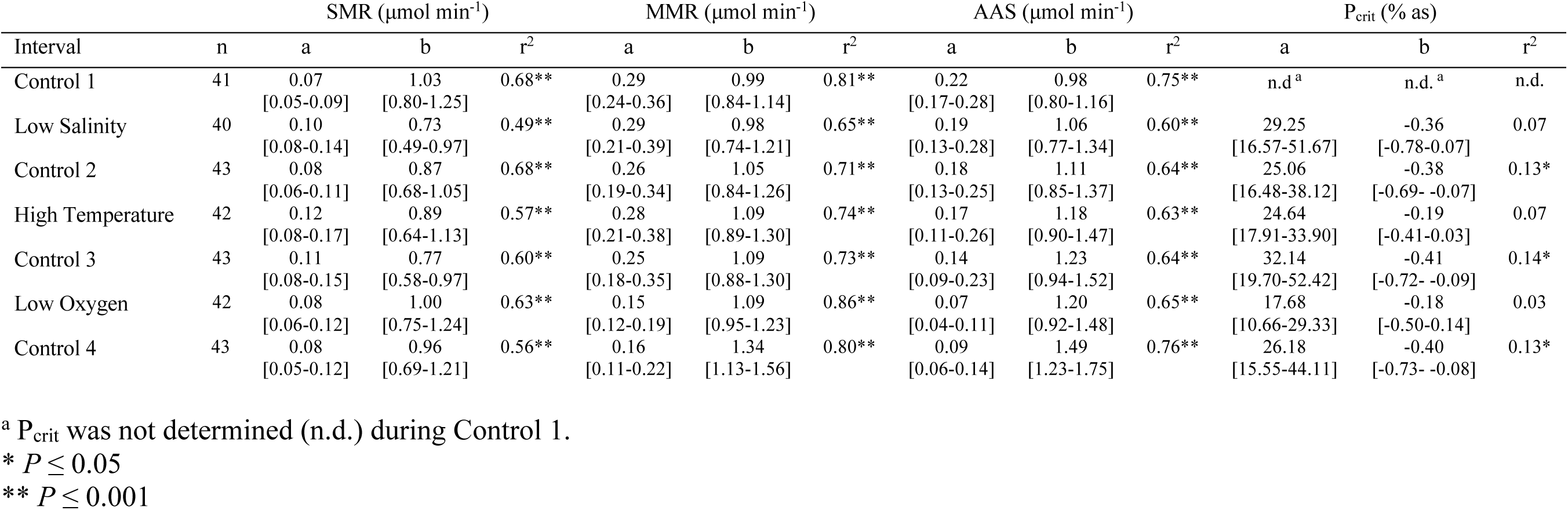
Scaling relationships for aerobic metabolic traits of *F. grandis* during laboratory acclimation. Coefficients were calculated for y = aM^b^, where y is the response variable, M is mass in g, and a and b are constants, shown with 95% confidence intervals in brackets. r^2^ values are for log-log regressions of mass versus the respective response variable.

| Interval | n | SMR ($\mu\text{mol min}^{-1}$) | | | MMR ($\mu\text{mol min}^{-1}$) | | | AAS ($\mu\text{mol min}^{-1}$) | | | $P_{\text{crit}}$ (% as) | | |
| --- | --- | --- | --- | --- | --- | --- | --- | --- | --- | --- | --- | --- | --- |
| | | a | b | $r^2$ | a | b | $r^2$ | a | b | $r^2$ | a | b | $r^2$ |
| Control 1 | 41 | 0.07<br>[0.05-0.09] | 1.03<br>[0.80-1.25] | 0.68** | 0.29<br>[0.24-0.36] | 0.99<br>[0.84-1.14] | 0.81** | 0.22<br>[0.17-0.28] | 0.98<br>[0.80-1.16] | 0.75** | n.d.<sup>a</sup> | n.d.<sup>a</sup> | n.d. |
| Low Salinity | 40 | 0.10<br>[0.08-0.14] | 0.73<br>[0.49-0.97] | 0.49** | 0.29<br>[0.21-0.39] | 0.98<br>[0.74-1.21] | 0.65** | 0.19<br>[0.13-0.28] | 1.06<br>[0.77-1.34] | 0.60** | 29.25<br>[16.57-51.67] | -0.36<br>[-0.78-0.07] | 0.07 |
| Control 2 | 43 | 0.08<br>[0.06-0.11] | 0.87<br>[0.68-1.05] | 0.68** | 0.26<br>[0.19-0.34] | 1.05<br>[0.84-1.26] | 0.71** | 0.18<br>[0.13-0.25] | 1.11<br>[0.85-1.37] | 0.64** | 25.06<br>[16.48-38.12] | -0.38<br>[-0.69- -0.07] | 0.13* |
| High Temperature | 42 | 0.12<br>[0.08-0.17] | 0.89<br>[0.64-1.13] | 0.57** | 0.28<br>[0.21-0.38] | 1.09<br>[0.89-1.30] | 0.74** | 0.17<br>[0.11-0.26] | 1.18<br>[0.90-1.47] | 0.63** | 24.64<br>[17.91-33.90] | -0.19<br>[-0.41-0.03] | 0.07 |
| Control 3 | 43 | 0.11<br>[0.08-0.15] | 0.77<br>[0.58-0.97] | 0.60** | 0.25<br>[0.18-0.35] | 1.09<br>[0.88-1.30] | 0.73** | 0.14<br>[0.09-0.23] | 1.23<br>[0.94-1.52] | 0.64** | 32.14<br>[19.70-52.42] | -0.41<br>[-0.72- -0.09] | 0.14* |
| Low Oxygen | 42 | 0.08<br>[0.06-0.12] | 1.00<br>[0.75-1.24] | 0.63** | 0.15<br>[0.12-0.19] | 1.09<br>[0.95-1.23] | 0.86** | 0.07<br>[0.04-0.11] | 1.20<br>[0.92-1.48] | 0.65** | 17.68<br>[10.66-29.33] | -0.18<br>[-0.50-0.14] | 0.03 |
| Control 4 | 43 | 0.08<br>[0.05-0.12] | 0.96<br>[0.69-1.21] | 0.56** | 0.16<br>[0.11-0.22] | 1.34<br>[1.13-1.56] | 0.80** | 0.09<br>[0.06-0.14] | 1.49<br>[1.23-1.75] | 0.76** | 26.18<br>[15.55-44.11] | -0.40<br>[-0.73- -0.08] | 0.13* |

<sup>a</sup> $P_{\text{crit}}$ was not determined (n.d.) during Control 1.
\* $P \leq 0.05$
\*\* $P \leq 0.001$

### Acclimation effects on aerobic metabolism

SMR, MMR, AAS, and P_crit_ were determined for *F. grandis* after acclimation to low salinity, high temperature, or low oxygen. Because these metabolic traits are affected by body mass, and because body mass increased over the course of the experiment (Table S3), mass-adjusted values for each variable were determined from log-log relationships with body mass and presented for a fish of average mass (4.39 g) for visualization purposes (Table 2; Fig. 2). Linear mixed models assessed the effects of fixed factors (sex, collection site, acclimation condition, and experimental interval), while accounting for body mass (log_10_ transformed) and individual (as a random factor) (Table 3).

**Table 2:** Mass-corrected aerobic metabolic traits (mean ± S.D.) of *F. grandis* during laboratory acclimation to changes in salinity, temperature, and dissolved oxygen. Values of SMR, MMR, AAS, and P_crit_ were determined for a fish of 4.39 g (average ass) based upon log-log relationship of each variable and body mass. The range is shown in parentheses.

| Interval | n | SMR<br>($\mu\text{mol min}^{-1}$) | MMR<br>($\mu\text{mol min}^{-1}$) | AAS<br>($\mu\text{mol min}^{-1}$) | $P_{crit}$<br>(% as) |
| --- | --- | --- | --- | --- | --- |
| Control 1 | 41 | 0.30 ± 0.05<br>(0.22 – 0.40) | 1.21 ± 0.13<br>(0.92 – 1.47) | 0.91 ± 0.11<br>(0.62 – 1.12) | n.d.<sup>a</sup> |
| Low Salinity | 40 | 0.31 ± 0.06<br>(0.21 – 0.43) | 1.19 ± 0.20<br>(0.73 – 1.75) | 0.88 ± 0.19<br>(0.50 – 1.41) | 18.2 ± 6.0*<br>(10.8 – 32.3) |
| Control 2 | 43 | 0.30 ± 0.04<br>(0.23 – 0.40) | 1.19 ± 0.20<br>(0.88 – 1.60) | 0.89 ± 0.18<br>(0.61 – 1.35) | 15.0 ± 4.6<br>(11.1 – 33.3) |
| High Temperature | 42 | 0.42 ± 0.07*<br>(0.30 – 0.60) | 1.40 ± 0.21*<br>(1.03 – 1.89) | 0.97 ± 0.21*<br>(0.65 – 1.38) | 19.0 ± 3.0*<br>(13.4 – 26.3) |
| Control 3 | 43 | 0.35 ± 0.05<br>(0.26 – 0.48) | 1.25 ± 0.18<br>(0.84 – 1.53) | 0.90 ± 0.18<br>(0.50 – 1.18) | 18.3 ± 4.4<br>(12.4 – 32.3) |
| Low Oxygen | 42 | 0.36 ± 0.06*<br>(0.23 – 0.56) | 0.76 ± 0.08*<br>(0.56 – 0.92) | 0.40 ± 0.08*<br>(0.20 – 0.56) | 14.4 ± 4.7*<br>(9.5 – 36.9) |
| Control 4 | 43 | 0.31 ± 0.06<br>(0.20 – 0.47) | 1.19 ± 0.20<br>(0.75 – 1.84) | 0.89 ± 0.19<br>(0.45 – 1.54) | 15.0 ± 4.2<br>(10.6 – 31.0) |

<sup>a</sup> $P_{crit}$ was not determined (n.d.) during Control 1.
\* Variable was significantly affected by acclimation condition (LMM, Table 3).

**Table 3:**
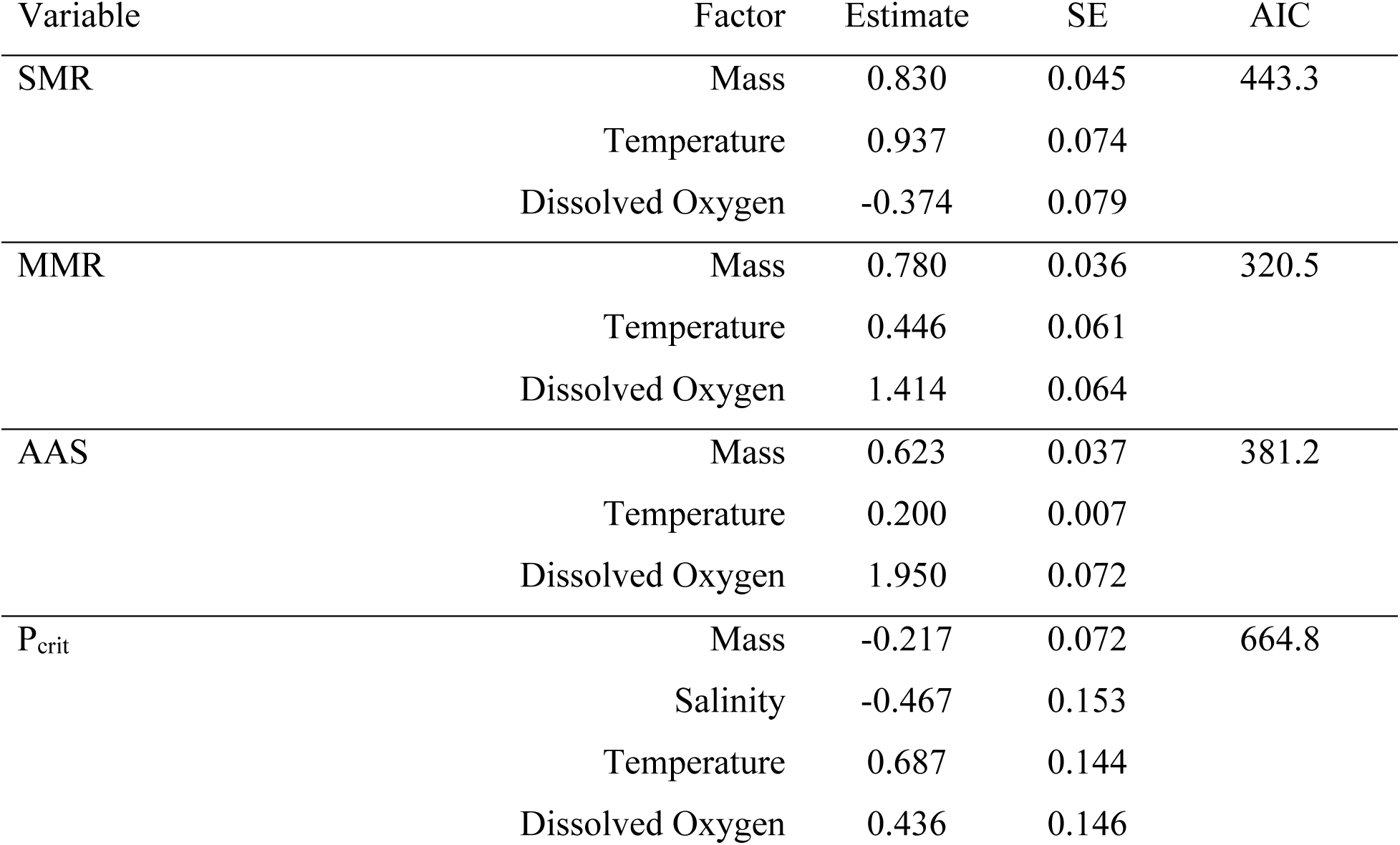
Factors influencing aerobic metabolic variables of *F. grandis* during laboratory acclimation to changes in salinity, temperature, and dissolved oxygen. Univariate LMMs were fit for SMR, MMR, AAS, and P_crit_ with collection site, sex, mass, salinity, temperature, dissolved oxygen, and experimental interval as fixed factors and individual ID as a random (intercept) factor. Response variables and body mass were log_10_ transformed and z-transformed. The minimum adequate model for each response variable is presented with corresponding AIC (Akaike Information Criterion).

**Figure 2:**
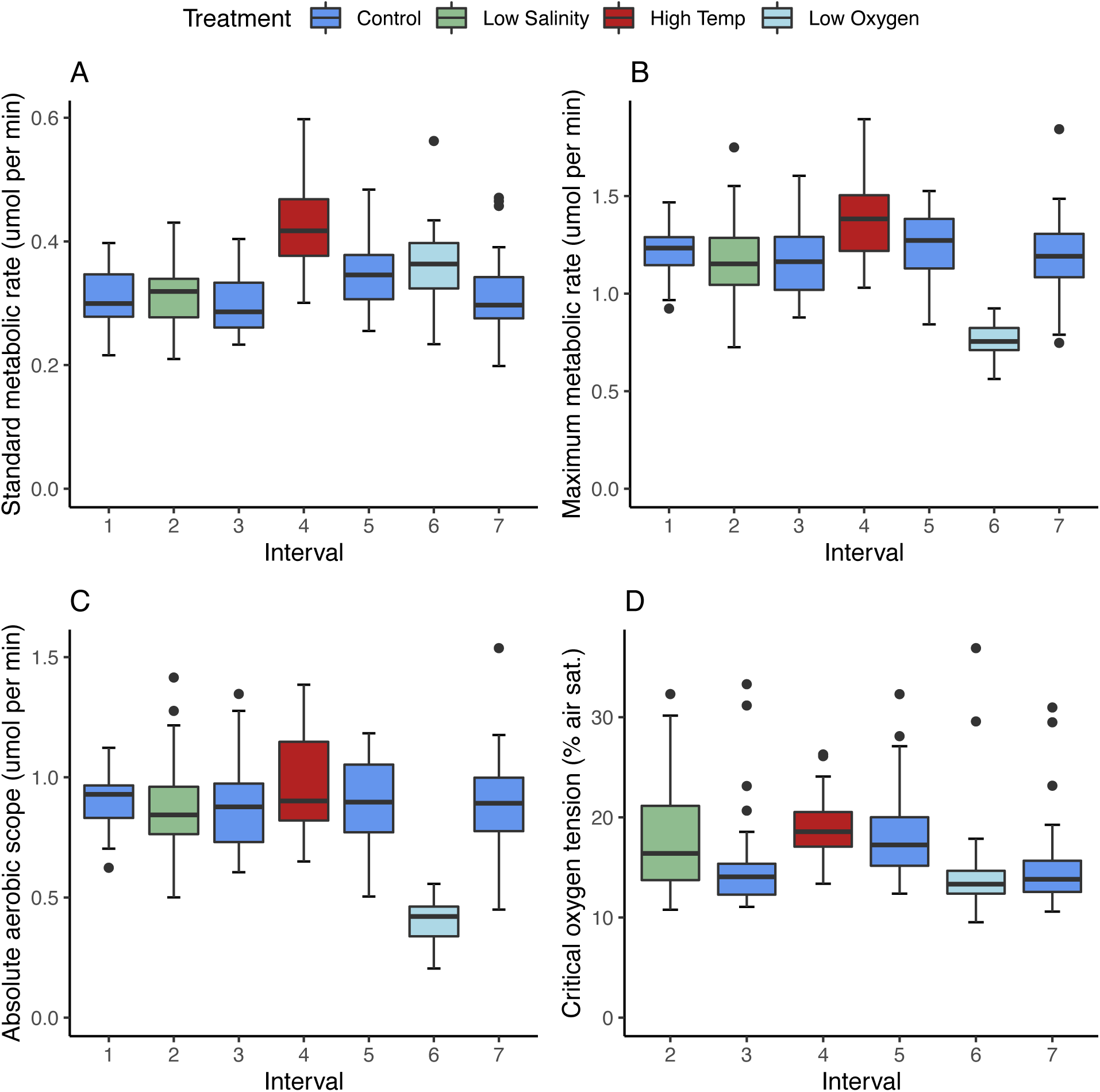
Aerobic metabolic traits of *F. grandis* during laboratory acclimation to low salinity (green), high temperature (red), and low oxygen (light blue). Measurements were made under control conditions (blue) before and after every acclimation interval. SMR (A); MMR (B); AAS (C); P_crit_ (D). All variables have been standardized for a fish of 4.39 g (the overall average mass). Box and whisker graphs show medians (centre line), upper and lower quartiles (box), and total data range (whiskers) after removing outliers (black dots).

SMR of *F. grandis* was not affected by acclimation to low salinity, but it was significantly elevated after acclimation to high temperature or low oxygen (Fig. 2A, Tables 2, 3). At 32°C, SMR was 33% higher than the average value recorded under control conditions (25°C), while at 30% a.s., SMR was about 14% higher than normoxic controls (Table 2). MMR was similarly unaffected by acclimation to low salinity and increased significantly after acclimation to high temperature (by 15% over control conditions); however, unlike SMR, MMR was dramatically suppressed (by 37%) at low oxygen (Fig. 2B, Tables 2, 3). Changes in AAS, mirrored those of MMR, being unaffected by acclimation to low salinity, increasing after acclimation to high temperature (but only by 8%), and decreasing after acclimation to low oxygen (by 55%) (Fig. 2C, Tables 2, 3). Changes in P_crit_ were generally small (Fig. 2D, Table 2) but significant (Table 3). P_crit_ was modestly elevated after acclimation to low salinity or high temperature, while it decreased after acclimation to low oxygen. The effects of high temperature and low oxygen on P_crit_ appeared to persist through the control interval that followed the respective acclimation period (Fig. 2D).

### Lack of collection site, sex, and interval effects

Although the two collection sites differed in DO profiles, both annually (Fig. 1) and at the time of fish collection (Table S1), collection site failed to explain significant variation in any of the measured traits related to aerobic metabolism. Moreover, the interaction between collection site and low oxygen treatment did not explain significant variation in any variable, indicating that fish from these two sites responded similarly to low oxygen acclimation. Additionally, there was no difference between sexes for any trait related to aerobic metabolism measured here. Finally, experimental interval did not explain significant variation in any variable, demonstrating that there was no temporal effect on traits related to aerobic metabolism over the 7-month laboratory experiment.

### Repeatability of aerobic metabolism

Pearson’s product moment correlation coefficients (r) were calculated to compare each response variable measured among all individuals between all possible pairs of intervals. In general, values of r were positive, but varied in magnitude and statistical significance among metabolic traits (Table S5). Thirteen of 21 pairwise comparisons between trials of SMR were significant; 16 were significant for MMR; and 11 were significant for AAS. Only three out of 15 comparisons were significant for P_crit_. There were no obvious patterns indicating that certain acclimation conditions were more or less likely to be correlated with each other or with control intervals.

The adjusted repeatability (R_adj_) evaluates trait consistency across the entire experiment, rather than between pairs of intervals, and accounts for main effects on trait values. To determine whether acclimation influenced the repeatability of the metabolic traits in question, R_adj_ was calculated two ways: first, R_adj_ was determined using data collected only during the control intervals, and second, R_adj_ was calculated over all intervals including acclimation treatments (Table 4). For control conditions, R_adj_ varied from 0.11 for P_crit_ to 0.36 for MMR. When including acclimation intervals, R_adj_ varied from 0.16 for P_crit_ to 0.37 for SMR. All values of R_adj_ were significantly different from zero, indicating that all traits were repeatable over the 7-month experiment, although there was a trend of lower R_adj_ for P_crit_ compared to the other variables. This result is consistent with the lower number of significant pairwise correlations for P_crit_ than for SMR, MMR, and AAS (Table S5). Moreover, values of R_adj_ were not materially affected when acclimation intervals were included in its calculation, suggesting that the repeatability of these traits is not influenced by acclimation to different conditions.

**Table 4:** Adjusted repeatabilities (R_adj_) of SMR, MMR, AAS, and P_crit_ of *F. grandis* measured during long-term laboratory maintenance. Values were determined for control intervals only, as well as across all intervals including acclimation to changes in salinity, temperature, and dissolved oxygen.

| Variable | Control Intervals |  |  |  | All Intervals |  |  |  |
| --- | --- | --- | --- | --- | --- | --- | --- | --- |
| | $R_{adj}$ | SE | 95% CI | p | $R_{adj}$ | SE | 95% CI | p |
| SMR | 0.35 | 0.09 | 0.18-0.51 | <0.001 | 0.37 | 0.07 | 0.23-0.51 | <0.001 |
| MMR | 0.36 | 0.08 | 0.18-0.51 | <0.001 | 0.35 | 0.07 | 0.21-0.48 | <0.001 |
| AAS | 0.32 | 0.09 | 0.14-0.47 | <0.001 | 0.27 | 0.07 | 0.14-0.40 | <0.001 |
| $P_{crit}$ | 0.11 | 0.06 | 0.01-0.23 | 0.007 | 0.16 | 0.07 | 0.05-0.30 | <0.001 |

### Phenotypic correlations between aerobic metabolic traits

Phenotypic correlations between pairs of metabolic variables were determined and partitioned into among-individual and within-individual correlations (Table 5, Fig. 3). There was a positive phenotypic correlation between SMR and MMR. This correlation arose from significant among-individual and within-individual correlations. Likewise, there was a positive phenotypic correlation between MMR and AAS, that was attributed to significant among-individual and within-individual correlations. On the other hand, there was no correlation between SMR and AAS at any level. The relationship of AAS with MMR, but not SMR, shows that the major factor determining AAS is MMR. There was a positive phenotypic correlation between SMR and P_crit_. While the magnitudes of the among-individual and within-individual correlations were similar, only the within-individual correlation was significant. Thus, for a given individual at a given time point, when SMR was high, so was P_crit_, and *vice versa*. This stands to reason because the determination of P_crit_ depends upon SMR (see Materials and Methods). Finally, there was a negative phenotypic correlation between AAS and P_crit_, which was attributed to a significant, negative within-individual correlation. Thus, in a trial when a given individual had a high P_crit_, it had a relatively low AAS.

**Table 5:** Phenotypic (r_p_), among-individual (r_ind_), and within-individual (r_e_) correlations between aerobic metabolic traits of *F. grandis*. The highest posterior distribution (HPD) interval was calculated for each estimate as a measure of credibility. When the HPD does not overlap zero, the correlation is significant (bold type).

| Correlation | $r_p$ | HPD Interval | $r_{ind}$ | HPD Interval | $r_e$ | HPD Interval |
| --- | --- | --- | --- | --- | --- | --- |
| SMR vs MMR | **0.24** | **0.17 – 0.32** | **0.68** | **0.37 – 0.88** | **0.28** | **0.17 – 0.40** |
| SMR vs AAS | -0.04 | -0.12 – 0.05 | 0.14 | -0.20 – 0.49 | -0.09 | -0.20 – 0.04 |
| SMR vs $P_{crit}$ | **0.31** | **0.19 – 0.40** | 0.32 | -0.03 – 0.66 | **0.36** | **0.24 – 0.48** |
| MMR vs AAS | **0.57** | **0.52 – 0.66** | **0.64** | **0.43 – 0.82** | **0.87** | **0.84 – 0.90** |
| MMR vs $P_{crit}$ | 0.02 | -0.10 – 0.11 | -0.13 | -0.24 – 0.50 | -0.01 | -0.17 – 0.11 |
| AAS vs $P_{crit}$ | **-0.11** | **-0.22 – -0.01** | 0.09 | -0.34 – 0.41 | **-0.20** | **-0.32 – -0.06** |

**Figure 3:**
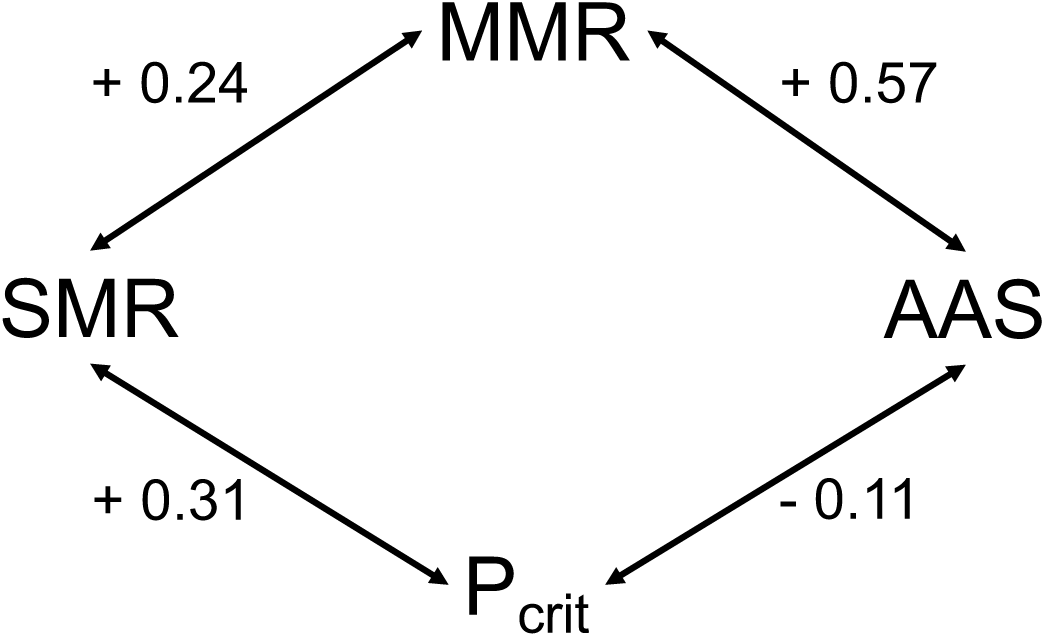
Phenotypic correlations (r_p_) between aerobic metabolic traits (SMR, MMR, AAS, and P_crit_) in *F. grandis*. For among-individual (r_ind_), and within-individual (r_e_) correlations and credibility statistics, see Table 5.

## DISCUSSION

### Mass effects on aerobic metabolism

In this study, all metabolic variables were significantly influenced by body mass. For SMR, the scaling coefficients were similar to those found previously in this species (b=0.79; Reemeyer et al., 2019) and other teleosts (reviewed in Jerde et al., 2019). In a recent meta-analysis of the relationship between SMR and body mass in fishes, Jerde et al (2019) provided strong evidence of an intra-specific mass scaling exponent near 0.89, aligning closely with the values reported here (Table 1). In the present study, the scaling coefficients for MMR and AAS were slightly higher than those calculated for SMR (Table 1), supporting the suggestion that MMR often scales isometrically with mass (reviewed in Glazier, 2009). On the other hand, the relationship between body mass and P_crit_ was negative. If P_crit_ is an index of hypoxia tolerance (Speers-Roach et al., 2013; Rogers et al., 2016; Regan et al., 2019; Wood, 2018), this result suggests that larger individuals are more tolerant of hypoxia than smaller individuals. Over a 3-fold range of body masses, and using an average value of b = -0.32, the largest individual would have a P_crit_ 30% lower than the smallest fish. In other words, the SMR of the larger fish would not be limited until oxygen dropped to values considerably lower than those that limit SMR of the smaller fish. A previous study in *F. grandis* found a similar result (Everett and Crawford, 2009), however other studies have found no effect of body mass on P_crit_ in *F. grandis* (Virani and Rees, 2000) or in *F. heteroclitus* (Borowiec et al., 2015; McBryan et al., 2016). Among other fishes, the relationship between P_crit_ and body mass is extremely variable, ranging from being positively related (Pan et al., 2016), to unrelated (Nilsson and Östlund-Nilsson, 2008; Timmerman and Chapman, 2004; Verheyen et al., 1994), to negatively related (Sloman et al., 2006; Perna and Fernandes, 1996; current study). While this diversity in scaling of P_crit_ may reflect real differences among species, it might also arise from different experimental and analytical methods used to determine P_crit_, highlighting the need for standardization in this area (Reemeyer and Rees, 2019; Regan et al., 2019; Wood, 2018).

### Acclimation effects on aerobic metabolism

Acclimation of *F. grandis* to low salinity brought about a small, but significant, increase in P_crit_, consistent with predictions based upon the osmoregulatory compromise (Sardella and Brauner, 2007). Giacomin et al. (2019) recently found higher P_crit_, lower gill surface area, and larger interlamellar cell masses in *F. heteroclitus* acclimated to freshwater (0 salinity) versus 11 and 35 salinity. Acclimation of *F. grandis* to low salinity, however, did not alter SMR, MMR, or AAS. In contrast, acclimation of *F. heteroclitus* to fresh water (0.3 salinity) significantly decreased factorial aerobic scope (FAS), calculated as MMR divided by routine metabolic rate (RMR) (Brennan et al., 2016). This effect was due to a trend toward lower in MMR of *F. heteroclitus* at low salinity (p = 0.06). This difference in response to low salinity between closely related species might be due to the use of FAS by Brennan et al. (2016) compared to AAS here. While both have their merits, AAS reflects the actual energy available to the organism to support activities beyond maintenance and, as such, has been argued to be more ecologically relevant (Clark et al., 2013). Results of the present study suggest that low salinity may hinder the ability to extract oxygen as DO drops (i.e., increase P_crit_), without altering the rate of oxygen uptake when oxygen is plentiful (SMR, MMR, and AAS were all determined at normoxia).

Acclimation of *F. grandis* to elevated temperature resulted in significant increases in all aerobic metabolic variables measured here. Based upon average values determined under control conditions (25°C), the increases in SMR, MMR, and AAS at 32°C correspond to Q_10_ values of 1.5, 1.2, and 1.1, respectively. While these values are lower than the range of 2 to 3 generally seen for aerobic metabolism of other fishes (Clarke and Johnston, 1999), they are consistent with the low temperature sensitivity of aerobic metabolism reported for *F. heteroclitus* acclimated to similar temperatures (Targett, 1978; Healy and Schulte, 2012). Specifically, Healy and Schulte (2012) showed that (RMR), MMR, and AAS sharply increased with an increase in acclimation temperature from 5 to 25°C, but then plateaued or decreased at higher acclimation temperatures (30 and 33°C). Taken together, these observations suggest that aerobic metabolism in these species is only moderately affected by temperature over the range studied here: The “oxygen and capacity limited thermal tolerance” theory suggests that MMR is more limited by temperature than SMR, leading to a diminution of AAS (Pörtner 2010). The above Q_10_ values suggest that SMR is more temperature-dependent than MMR, although this comparison belies the fact that the absolute increase in MMR was approximately twice that of SMR. In absolute terms, AAS increased at 32°C, albeit with a low Q_10_. Across this temperature range, therefore, aerobic metabolism by *F. grandis* does not appear to be limited. This conclusion is supported by the observation that growth rates were highest during acclimation to high temperature (Table S3), as well as with observations made in the field, where these fish were active at temperatures at or above 32°C.

P_crit_ values were also higher following acclimation of *F. grandis* to high temperature, suggesting that fish may be less tolerant to low oxygen as temperatures increase. Previous work showed that acute warming from of *F. heteroclitus* from 15 to 30°C led to a decrease in hypoxia tolerance when measured as time to loss of equilibrium (LOE) during exposure to severe hypoxia (2% a.s.; McBryan et al., 2016). Interestingly, acclimation to warm temperature partially reversed the negative effects of acute warming on LOE, a response that was correlated with an increase in gill surface area in warm-acclimated fish. Despite differences in experimental design (acclimation vs. acute exposure, P_crit_ vs. LOE), the current study and McBryan et al. (2016) both point to a decrease in hypoxia tolerance of *F. grandis* and *F. heteroclitus*, respectively, at higher temperatures. These results are consistent with observations and theoretical arguments made for other ectothermic species (reviewed in McBryan et al., 2013).

Acclimation of *F. grandis* to low oxygen led to decreases in MMR and AAS. This was expected due to the restriction of MMR, and hence AAS, at low DO, even at levels above P_crit_ (Richards, 2009; Rogers et al., 2016). Moreover, P_crit_ was also lowered, which supports previous work in *F. heteroclitus* (Borowiec et al., 2015) and in other fishes (reviewed in Rogers et al., 2016). This indicates that acclimation to low oxygen increases hypoxia tolerance, presumably through a variety of morphological, physiological, and biochemical adjustments (Richards, 2009; Sollid et al., 2003). Surprisingly, acclimation to hypoxia resulted in a slight increase in SMR. This result contrasts with Borowiec et al. (2015), who showed that acclimation of *F. heteroclitus* for 28 d to ∼24 % a.s. did not affect RMR measured in normoxia. The current observation of higher SMR under hypoxia might be explained, in part, by an experimental design in which MMR was measured prior to overnight determination of SMR. To induce MMR, fish were chased to exhaustion, a protocol that elicits anaerobic metabolism and lactate accumulation (Rees et al., 2009). Lactate clearance after exercise, either by oxidation or gluconeogenesis, results in an increase in oxygen-consumption, the well-described “excess post-exercise oxygen consumption” (Hill and Lupton, 1923; Scarabello et al., 1991; Wood, 1991). After a similar exercise protocol, blood lactate decreased to control values within 3 h under normoxia (Rees et al., 2009). This decrease would have occurred prior to the beginning of SMR measurements in normoxic exposures. Under hypoxia, though, EPOC probably lasts longer (Svendsen et al., 2012), and may have contributed to an elevation of SMR. An increased duration of increased oxygen consumption, after either exercise or the ingestion of food (specific dynamic action, Jobling, 1981; Chabot et al., 2016), could have important ecological implications, because it would contribute to a decrease in AAS and the energy available to perform other tasks. Indeed, AAS at low oxygen was dramatically reduced in the current study, being less than half the value measured during normoxic controls.

### Lack of collection site and sex effects on aerobic metabolism

Fish in this study were sampled from two sites within the GBNERR that differed in seasonal DO profiles, where one site (BH) experiences a higher frequency of hypoxia than the other (BC). It was hypothesized that due to these differences in DO, fish from BH may show fixed developmental or evolved differences in these metabolic variables (e.g. lower SMR and P_crit_) or the degree to which these variables responded to low oxygen acclimation. Previous work in *F. grandis* suggested that populations differed in M_O2_ at severe hypoxia (ca. 9% a.s.) but found no differences under normoxia nor differences in P_crit_ (Everett and Crawford, 2009). In sailfin mollies (*Poecilia latipinna*), a species in the same order (Cyprinidoniformes) and occurring in similar habitats as *F. grandis*, fish from a periodically hypoxic salt marsh have significantly lower P_crit_ and higher gill surface area than fish from a normoxic river site (Timmerman and Chapman, 2004). In the present study none of the response variables differed by collection site, nor did fish from the two sites differ in their response to acclimation low oxygen. There are at least three reasons why site-dependent differences were not observed in the present study. First, it is possible that there is no significant genetic differentiation between collection sites. These sites were separated by about 10 km, a distance much greater than the expected home range of this species (Nelson et al., 2014), but substantially less than that in studies where population differences were found [up to 650 km in Everett and Crawford (2009) and 78 km in Timmerman and Chapman (2004)]. Although migration of individual fish between sites probably does not occur, low rates of gene flow over several generations may be sufficient to overwhelm selection to local conditions (Slatkin, 1987). Second, it is possible that fish behaviourally avoid hypoxia and exploit microenvironments that have higher DO than those reflected in long-term data and point samples. For example, several species, including *F. grandis*, use aquatic surface respiration to improve oxygen uptake at low DO by ventilating their gills with surface waters that have higher levels of oxygen due to diffusion from the atmosphere (McKenzie and Chapman, 2009; Love and Rees, 2001; Rees and Matute, 2018). Third, the DO level chosen for acclimation might not have been low enough to elicit site-dependent differences. The DO level during acclimation (30% a.s.) was based upon data from field sites; however, this is above the P_crit_ of this species (Virani and Rees, 2000; present study). Acclimation to more severe hypoxia would be expected to recruit additional behavioural and physiological responses that might reveal site-dependent differences.

No metabolic trait measured here differed between male and female fish. This lack of sex effects contrasts the view that sex differences in reproductive investment and behaviour lead to differences in energy expenditure (Biro and Stamps, 2010). The lack of sex effects in this study might be attributed to the age of the fish used, which were probably young-of-the-year (Greeley and McGregor, 1983) and not reproductively active at the time of capture. In addition, fish in the current study were held at relatively high densities (∼60 fish per m^3^), which have been shown to reduce egg production in *F. grandis* (Chesser et al., 2019). On the other hand, over the course of laboratory maintenance, fish grew significantly and developed dimorphic coloration typical for sexually mature individuals of this species. Moreover, previous research on larger individuals of this species found no difference in SMR among males and females (Reemeyer et al., 2019). Thus, sex effects on aerobic metabolism in this species, if any, are small in magnitude.

### Repeatability of aerobic metabolism

All variables measured exhibited moderate repeatability over the course of the experiment. Values of R_adj_ ranged from 0.27 – 0.37 for SMR, MMR, and AAS, and fall within the range of values reported for aerobic metabolism (Nespolo and Franco, 2007; Norin and Malte, 2011; White et al., 2013). R_adj_ for P_crit_ was lower (0.11 – 0.16), suggesting that within-individual variation relative to among-individual variation was greater for this metric than for SMR, MMR, and AAS. Importantly, R_adj_ values reported here were determined over 7 months. Previous measurements of the repeatability of M_O2_ by *F. grandis* determined over shorter intervals have produced higher estimates of repeatability. Reemeyer et al. (2019) reported Pearson’s r ranging from 0.35 – 0.76 and an R_adj_ of 0.56 for SMR of *F. grandis* measured five times over six weeks. Virani and Rees (2000) reported a similarly high Pearson’s r for RMR (0.68) when measured twice within 6 weeks. Previous measurements of P_crit_ in *F. grandis* measured twice over 2 weeks resulted in a Pearson’s r of 0.74 (Reemeyer and Rees, 2019). The lower R_adj_ estimates found in the present study support the trend that repeatability of metabolic variables decreases over time as reported for other species (Norin and Malte, 2011; White et al., 2013). Nevertheless, the values R_adj_ reported here were all statistically greater than zero and suggest some degree of consistency among individuals with respect to aerobic metabolism.

While acclimation to altered salinity, temperature, and oxygen had significant effects on the magnitude of all metabolic variables measured here, R_adj_ estimates were virtually identical when determined on only control intervals and when calculated over the entire experiment, including acclimation intervals. Furthermore, pairwise correlations calculated between control intervals were similar to those calculated between control and acclimation intervals (Table S5). These observations suggest that the repeatability of these metabolic traits in *F. grandis* is not context dependent across this range of salinity, temperature, and DO. Auer et al. (2018) assessed the effect of temperature acclimation on repeatability of SMR, MMR, and AAS in juvenile brown trout (*Salmo trutta*) after serial acclimation to 10, 13, and 16°C. Although R_adj_ values were similar to those seen here (0.32 for SMR, 0.43 for MMR, and 0.42 for AAS), the R_adj_ of MMR and AAS, but not SMR, decreased with warming. For MMR and AAS, therefore, Auer et al (2018) provide some support for context-dependency of repeatability. In that study, fish were measured once at each temperature in the same order without a common control treatment between or after temperature acclimation; thus, it is possible that the lower R_adj_ determined at high temperatures were due, in part, by a time-dependent decrease in repeatability.

### Phenotypic correlations between aerobic metabolic traits

Because SMR, MMR, AAS, and P_crit_ were measured in a given respirometric trial and trials were repeated over time, it was possible to calculate phenotypic correlations between pairs of traits and partitioning these correlations into among-individual and within-individual correlations (Dingemanse and Dotchermann, 2012). The aerobic capacity model predicts a positive correlation between SMR and MMR (Bennett and Ruben, 1979; Hayes and Garland, 1995), however support for this relationship is mixed. A recent meta-analysis confirmed a positive relationship between SMR and MMR when comparing across species but failed to support a relationship between SMR and MMR within species (Auer et al., 2017). Similarly, Killen et al. (2016b) found that among species of teleost fishes, the correlation between RMR and MMR was strongly positive, but within species, the same correlation varied greatly. The present study provides strong evidence for a positive phenotypic correlation between SMR and MMR within a species. Moreover, this correlation can be attributed to significant correlations both among individuals and among repeated measures on the same individual. There was also a positive phenotypic correlation between MMR and AAS, which was due to significant among- and within-individual correlations. On the other hand, SMR and AAS were not correlated. Because AAS is calculated as the difference between MMR and SMR, the strong correlation between MMR and AAS highlights that variation in MMR is quantitatively more important in determining AAS than variation in SMR.

There was a positive phenotypic correlation between SMR and P_crit_. Although the among- and within-individual correlations were of similar magnitude, only the latter was statistically significant. Thus, during a given trial on a given individual, if SMR was elevated, P_crit_ was also elevated. While this result indicates that these variables may be linked by shared phenotypic plasticity, it would also arise from the method of determining P_crit_, which is directly dependent upon the magnitude of SMR (Reemeyer and Rees, 2019). Finally, there was a significant negative phenotypic correlation between P_crit_ and AAS, which was similarly attributed to a significant within-individual correlation. This would be predicted based upon the ways in which SMR is used to calculate P_crit_ and AAS. A lower SMR generally yields a lower P_crit_, but a higher AAS (depending upon variation in MMR, of course). Although this relationship can be explained by how these variables are calculated, it might have biological relevance. In particular, this result suggests that when a fish has a low SMR it simultaneously has a greater hypoxia tolerance (lower P_crit_) and a higher scope for activity (higher AAS).

These last relationships, which are driven primarily by within-individual variation, highlight the importance of repeated measures when examining the relationship between traits. Had these metabolic traits been measured only once among a group of individuals, significant correlations could have been attributed to among-individual differences, rather than covariation of these traits within an individual (Careau and Wilson, 2017). That conclusion would lead to experiments examining genetic or developmental factors that produced these correlations rather than an investigation of why those traits covary when measured multiple times for a given individual, perhaps due to shared plasticity or factors related to experimental design.

### Perspectives

Acclimation of SMR, MMR, AAS, and P_crit_ in *F. grandis* indicate plasticity of aerobic metabolism and may contribute to the broad environmental tolerances of this ecologically dominant estuarine species. It is possible that these tolerances, however, will be exceeded by changes in salinity, temperature, and DO that are more extreme, longer lasting, or contemporaneous. Predictions of future characteristics of estuaries are strongly influenced by local conditions (Wong et al., 2014). In the Southeast U.S.A., increased precipitation will likely decrease salinity, while simultaneously increasing nutrient input, eutrophication, and the incidence of aquatic hypoxia. Average and maximum water temperatures are projected to increase due to climate change (Pörtner et al., 2014). Thus, *F. grandis* will be challenged by increased oxygen demands at the same time as oxygen availability, directly or indirectly due to the osmoregulatory compromise, will limit their ability to meet those demands. Future studies on the effects of simultaneous variation in multiple abiotic factors will help elucidate the limits of resiliency of this and other estuarine organisms.

## List of symbols and abbreviations

a.s.: air saturation
AAS: Absolute aerobic scope
BC: Bayou Cumbest
BH: Bayou Heron
DO: dissolved oxygen
GBNERR: Grand Bay National Estuarine Research Reserve
M_b_: body mass
MMR: Maximum metabolic rate
M_O2_: rate of oxygen consumption
P_crit_: critical oxygen tension
RMR: routine metabolic rate
R_adj_: adjusted repeatability
r_p_: phenotypic correlation
r_e_: within-individual correlation
r_ind_: among-individual correlation
SMR: standard metabolic rate

## Acknowledgements

We thank Dr. Mark Woodrey and the staff of the Grand Bay National Research Estuarine Research Reserve for help in fish collection. We thank Mohammad Hamed and Bennett Price for assistance with fish care and Drs. Fernando Galvez and Simon Lailvaux for constructive comments on earlier versions of this manuscript.

## Author contributions

J.E.R. and B.B.R conceived and designed the methodology. J.E.R collected and analysed the data. J.E.R and B.B.R. interpreted the data. J.E.R drafted the manuscript. B.B.R edited the manuscript. Both authors contributed critically to the drafts and gave final approval for publication.

